# Exploring potential targets for quercetin treatment of colorectal cancer through network pharmacology and molecular docking technology

**DOI:** 10.1101/2024.07.29.605708

**Authors:** Yuanyuan Qian, Zhaojunli Wang, Jiancheng Ji, Wenlong Fu

## Abstract

**Objective:** To explore the potential targets of quercetin in the treatment of colorectal cancer through network pharmacology and molecular docking technology.

**Method:** Download the 3D structure of quercetin from Pubchem database, use PharmaMapper database for reverse docking, and screen protein targets based on zscore; GEO database screening for colorectal cancer datasets and screening for differentially expressed genes; Screening common genes for protein targets and differentially expressed genes using Wayne diagrams; Use DAVID database for GO and KEGG analysis. Use ChemDraw, 20.0 AutoDock, and Pymol for molecular docking.

**Result:** The results showed that 24 common genes and 13 signaling pathways were identified from the reverse docking data of quercetin and the GSE33113 gene chip data of colorectal cancer; Molecular docking results showed that quercetin showed good binding energy with 1ykc, 1k3y, 1kbq, 2bkz, 1xo2, 1og5, 2bsm.

**Conclusion:** Quercetin may exert its therapeutic effect on colorectal cancer through a multi target and multi channel mechanism.

## Introduction

Colorectal cancer is derived from epithelial tissue, occurs in the colon and rectum of a common form of gastrointestinal malignant tumor, morbidity after age 40 more[1], the morbidity after gastric cancer and esophageal cancer, serious impact on human health. With the improvement of our country people’s quality of life and eating habits changing, colon cancer incidence and mortality rate has arising trend[2-4], in one of the men and the elderly is a major population colorectal cancer[5].

Quercetin, also known as quercetin, is a kind of flavonol compound with a variety of biological activity[6], quercetin as antiapoptotic, antioxidants can protect to multiple organ damage, And in anti-cancer anti-inflammatory[7,8], antibacterial[9], antiviral [10], fall blood sugar, blood pressure [11], adjust the immune function and protect cardiovascular therapy effect[12].

Is based on the theory of systems biology and network pharmacology, network analysis of biological systems, selecting a specific signal node to multiple targets for drug molecular design of new subject. Network pharmacology emphasizes much way of signaling pathways regulate, to improve the therapeutic effect of drugs, reduce side effects, so as to improve the success rate of new drug clinical trials, save drug research and development costs. With the rise of network pharmacology, make its are widely applied to explore the traditional Chinese medicine and compound ingredients, targets, such as multiple pathways direction[13-15].

Molecular docking is refers to the receptor through energy, space match between matching and form compounds, and chemical properties and predict complex structure and affinity of a theoretical simulation method. In recent years, molecular docking methods have become an important technique in the field of computer-aided drug research[16,17].

In this study, network pharmacology and gene chip data were used to explore the potential targets of quercetin in the treatment of colorectal cancer, and molecular docking technology was used to explore the binding ability of quercetin to target proteins to verify the potential of quercetin in the treatment of colorectal cancer.

## 1. Website and software

### 1.1 Website

Pubchem database (https://pubchem.ncbi.nlm.nih.gov/);

PharmMapper database (http://www.lilab-ecust.cn/pharmmapper/);

Uniprot database (https://www.uniprot.org/);

DAVID database (https://david.ncifcrf.gov/home.jsp);

GEO database (https://www.ncbi.nlm.nih.gov/geo/);

Microscopic letter (http://www.bioinformatics.com.cn/);

Jvenn (https://jvenn.toulouse.inrae.fr/app/index.html).

PDB Protein database (https://www.rcsb.org/).

### 1.2 software

Cytoscape; ChemDraw 20.0; AutoDock; pymol; openBabel.

## 2. Methods

### 2.1 Quercetin 3D structure Download

Through the Pubchem database download quercetin 3 d structure, and through openBabel software will SDF files into mol2 files.

### 2.2 Reverse docking of quercetin

Upload quercetin mol2 files to reverse docking PharmMapper database, and according to the zscore is greater than 1 filter condition to screening of protein, and use Cytoscape draw quercetin - target protein (PDB ID) network diagram.

### 2.3 PDB ID converted to SYMBOL ID

Convert PDB ID to Uniprot Entry ID through Uniprot database, and then convert Uniprot Entry ID to SYMBOL ID through DAVID database.

### 2.4 screening of colorectal cancer datasets

The GEO database was used to screen the colorectal cancer datasets, and the datasets about colorectal cancer tissues and normal colon tissues were screened.

### 2.5 gene screening differences

According to genetic variations (2.4) selection datasetselection, and according to the |Log (FC)| > 1 and P values < 0.05 screening differences and map the volcano.

### 2.6 screening common genes

Through Jvenn website with Wayne figure screening (2.3) and (2.5) of common genes, and use Cytoscape draw quercetin - target protein (PDB ID) - genetic network diagram.

### 2.7 GO,KEGG analysis

Will choose common gene by DAVID database (2.6) to carry on the GO, KEGG analysis, and according to the P Value < 0.05 screening, will GO (BP_CC_MF) by microscopic map into a letter, by KEGG pathway analysis genetic screening. Cytoscape was used to draw the quercetin-target protein (PDB ID) -gene-pathway network diagram.

### 2.8 the protein file downloads

Download protein files in the PDB Protein database by target protein (PDB ID).

### 2.9 Molecular Docking

(1) Before treatment: quercetin Pubchem 3 d file imports ChemDraw 3 din the lowest energy; The target protein files were imported into pymol software for dehydration-hydrogen-deligandation to separate the monomer proteins.
(2) Molecular docking: use AutoDockTools to hydrogenation and charge the target protein; Hydrogenation of quercetin plus charge and Root; Molecular docking was performed using autogrid4 and autodock4.
(3) Visual processing: using openBabel software will PDBQT files into a PDB file, use pymol visualization processing software.

## 3. Result

### 3.1 Quercetin target screening

The quercetin 3D structure was downloaded from Pubchem database, reverse docking was performed from PharmMapper database, and the proteins were screened according to the zscore greater than 1 screening condition, and 93 target proteins were screened (Figure 1). Uniprot database and DAVID database are used to convert the PDB ID into the SYMBOL ID, get 96 SYMBOL ID.

**Figure 1.**
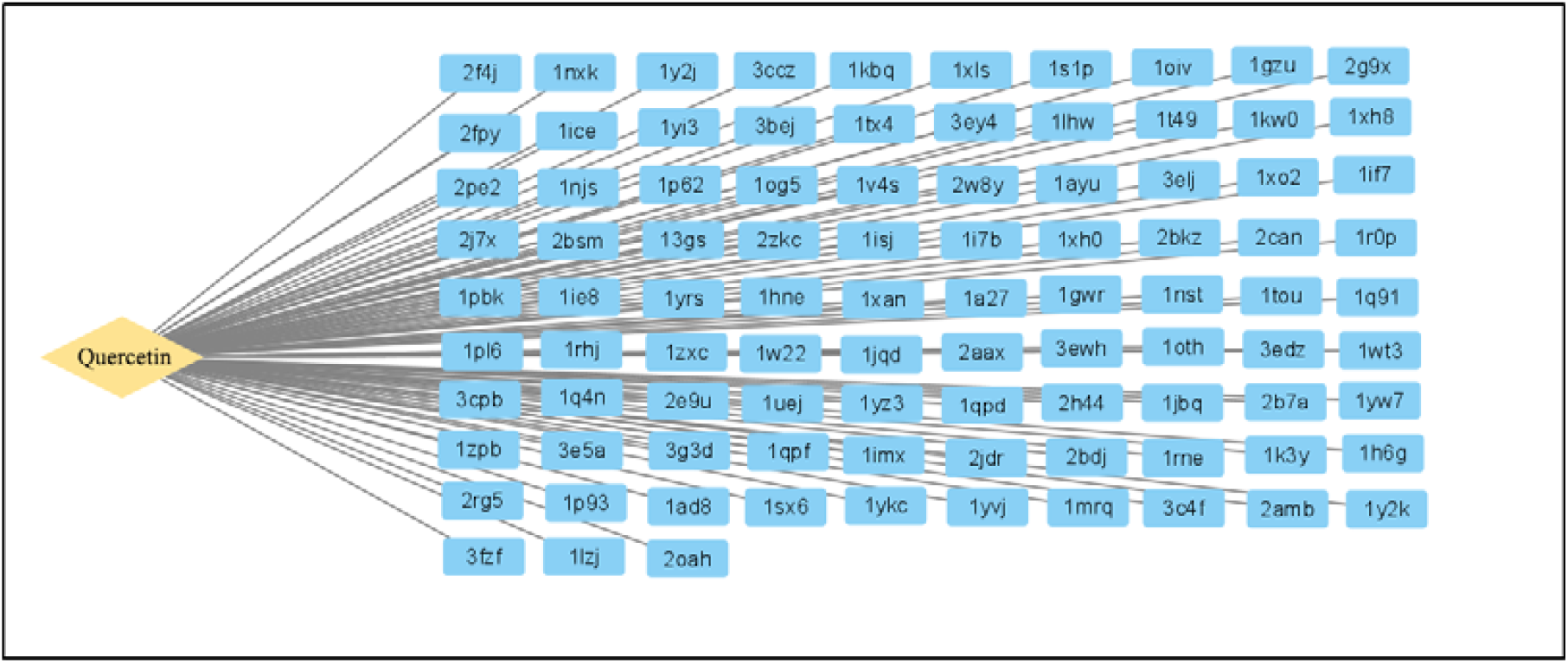
Quercetin-target protein (PDB ID) network diagram.

### 3.2 Targets associated with colorectal cancer

Screening colorectal cancer in the GEO database GSE33113 gene chip data and filter difference, after |Log (FC)| > 1 and P values < 0.05 screening, screening 3024 raised genetic genes and 1155 down-regulated (Figure 2), no corresponding gene were excluded and duplicate names were screened 2610 different genes.

**Figure 2.**
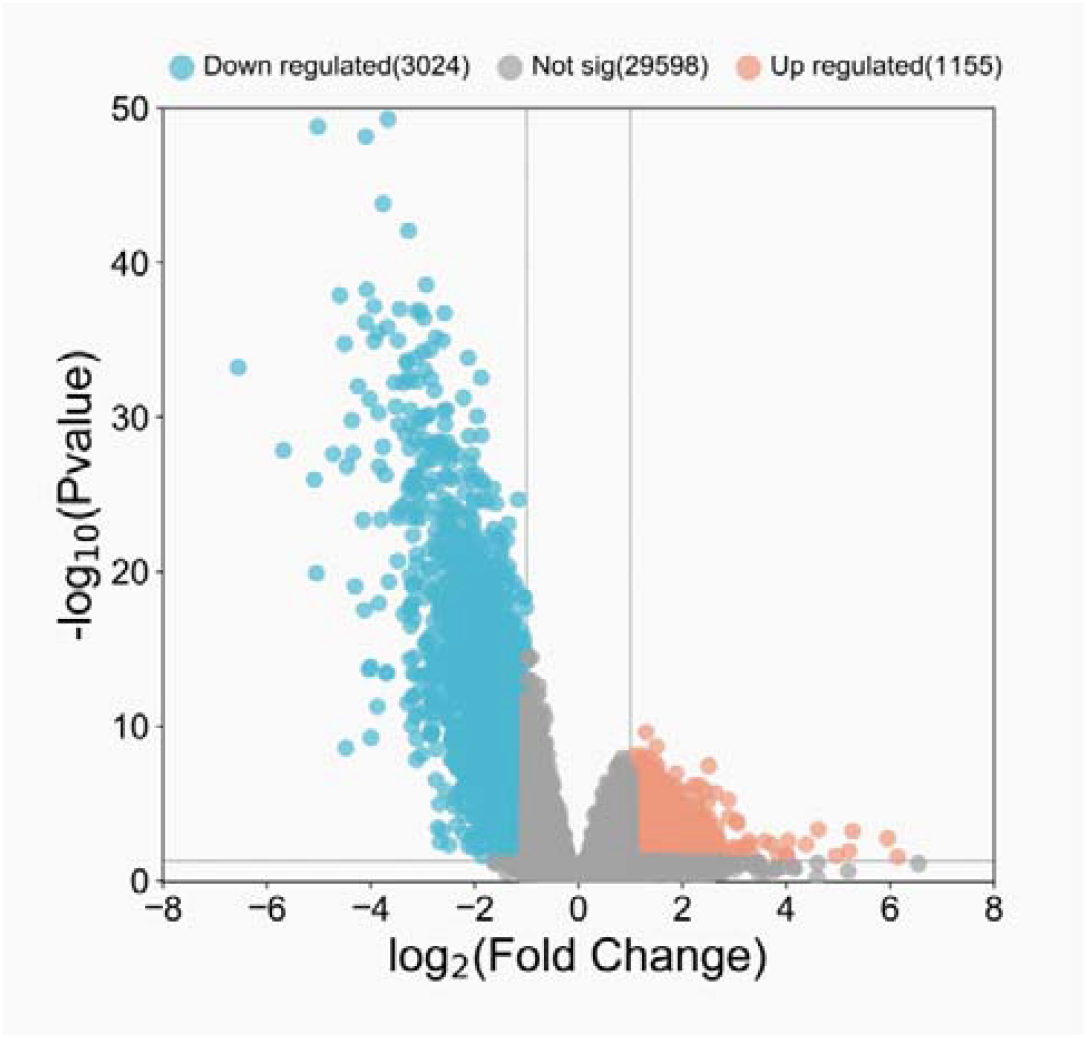
GSE33113 colorectal cancer gene chip volcanic figure.

### 3.3 Screening common genes

Use Wayne figure common genetic screening, screening 24 common gene (Figure 3), and use Cytoscape draw quercetin - target protein gene (PDB ID) - network diagram (Figure 4).

**Figure 3.**
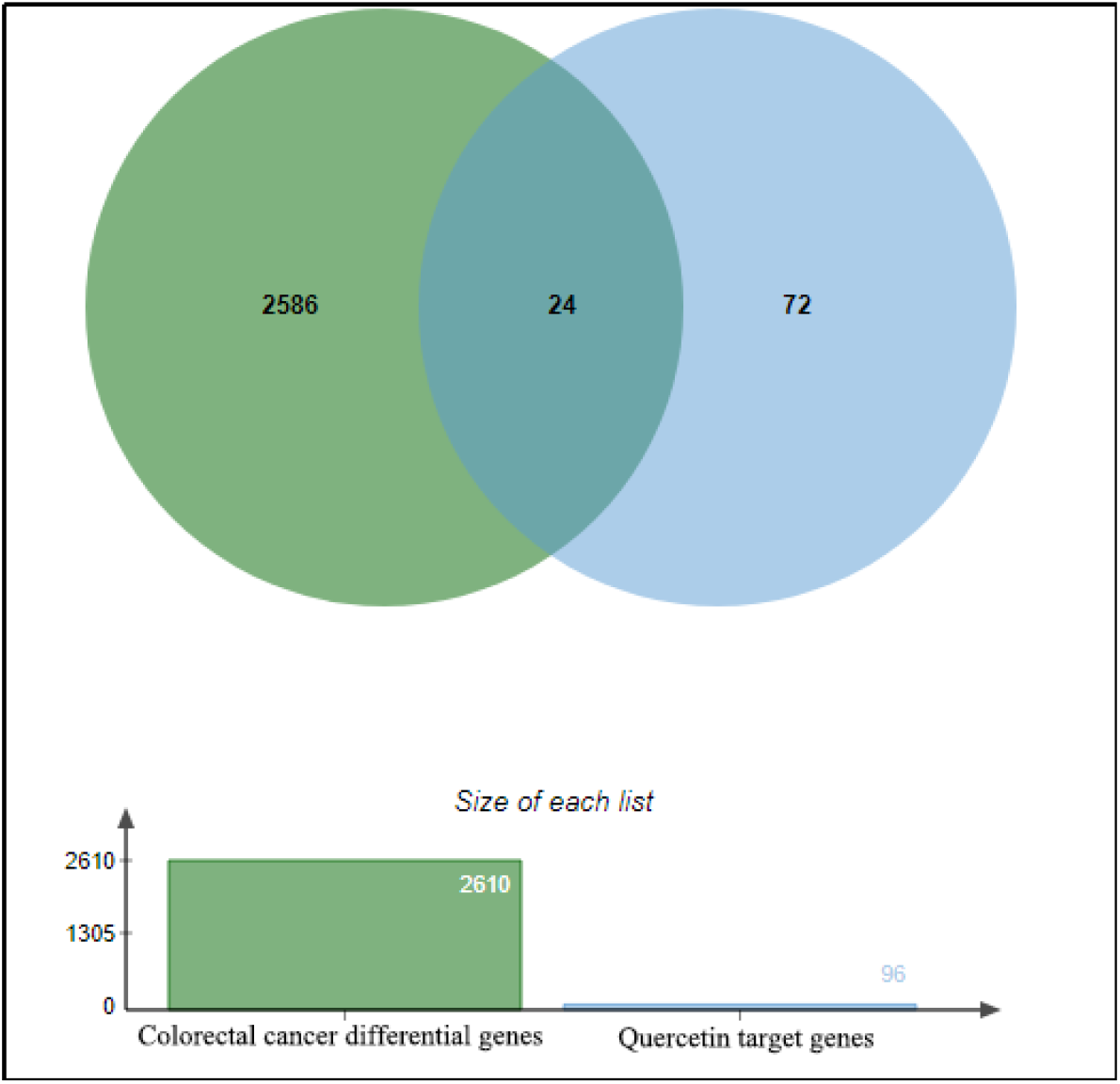
Differences between colorectal cancer genes and quercetin target genes Wayne figure.

**Figure 4.**
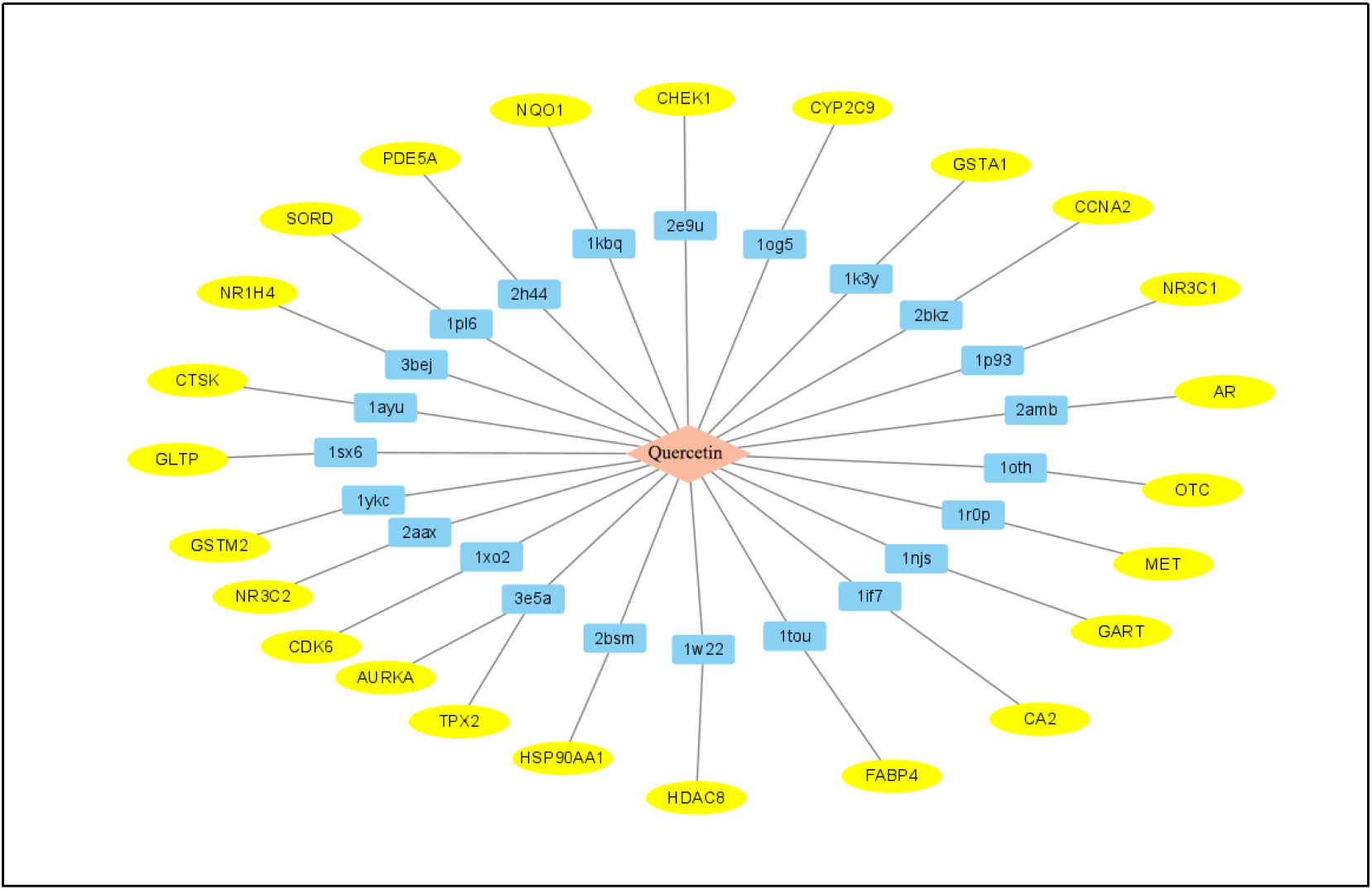
Quercetin - target protein (PDB ID) - genetic network diagram.

### 3.4 GO, KEGG analysis

Will choose the common genes into (3.3) to GO to DAVID database, KEGG analysis (Figure 5, Figure 6), according to the result of KEGG out seven target protein molecular docking, with Cytoscape drawing quercetin - target protein gene (PDB ID) - only strengthen network diagram (Figure 7).

**Figure 5.**
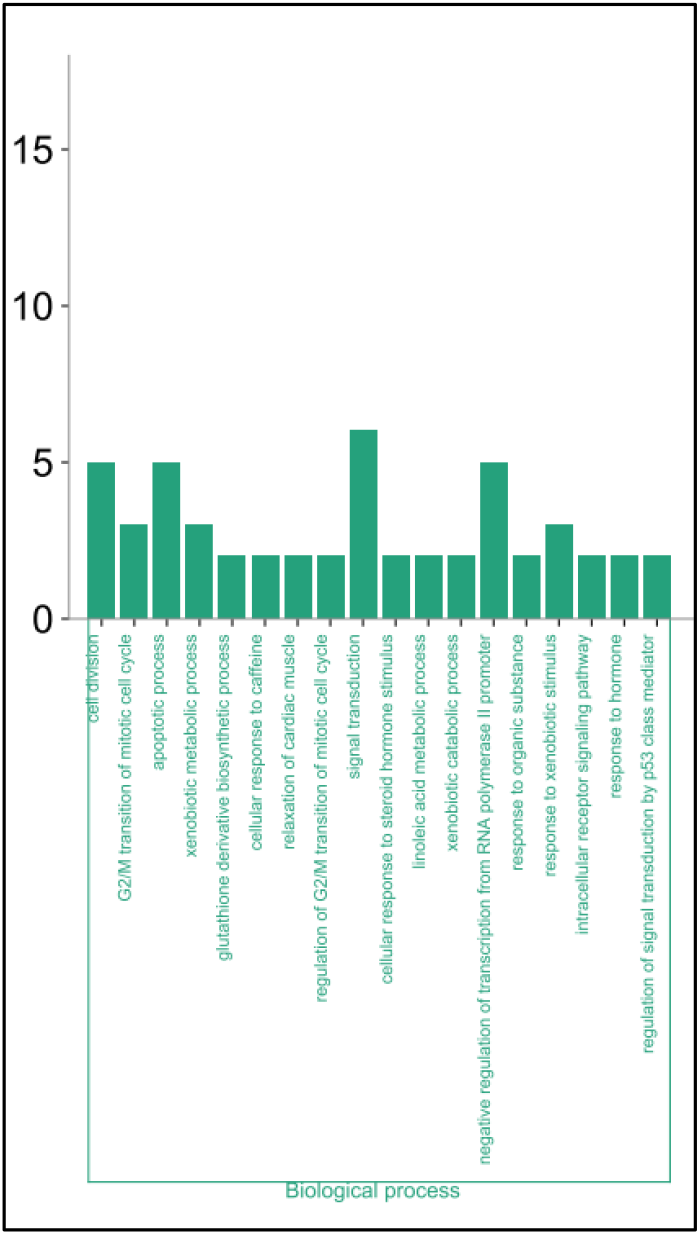
Visualization of GO BP results.

**Figure 6.**
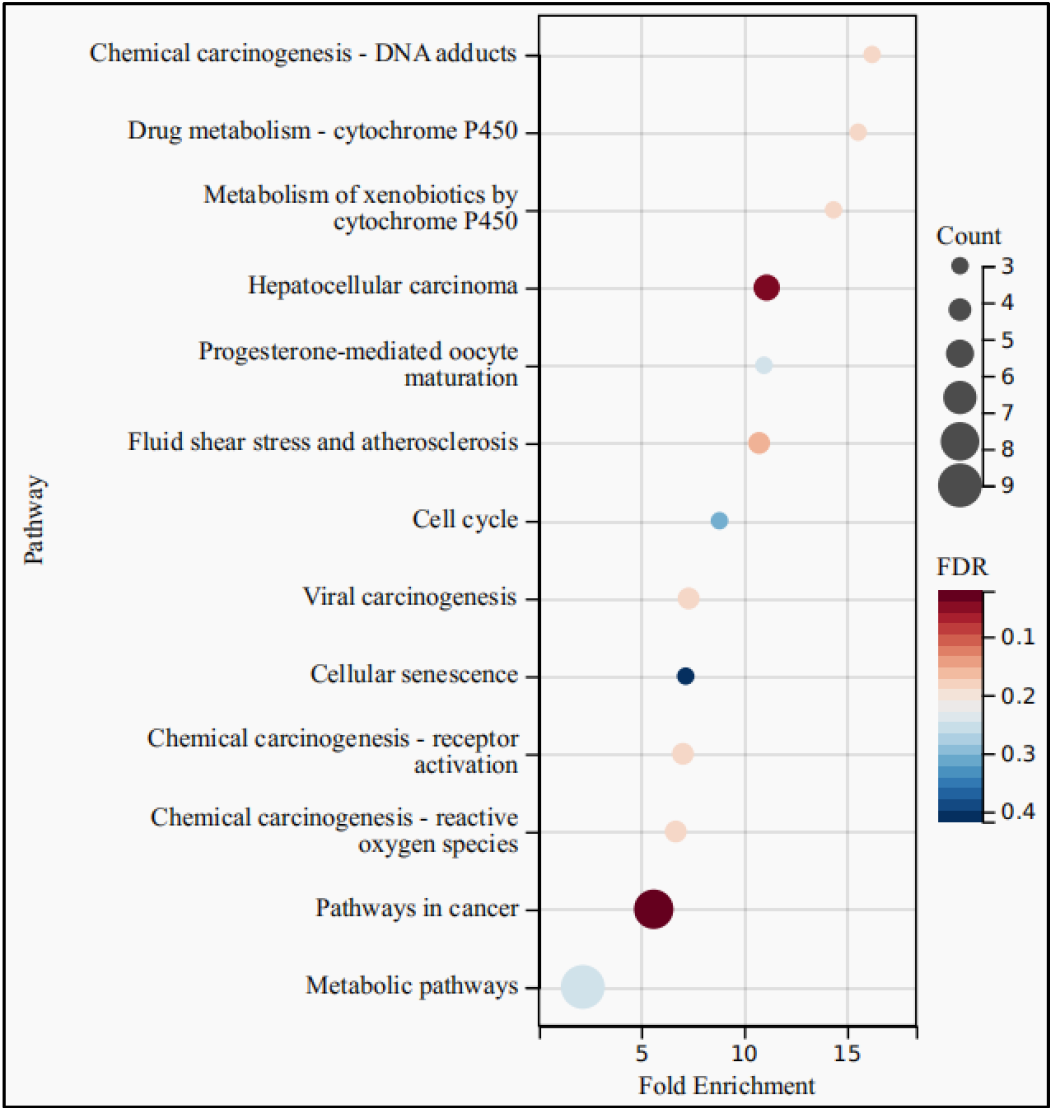
Visualization of KEGG results.

**Figure 7.**
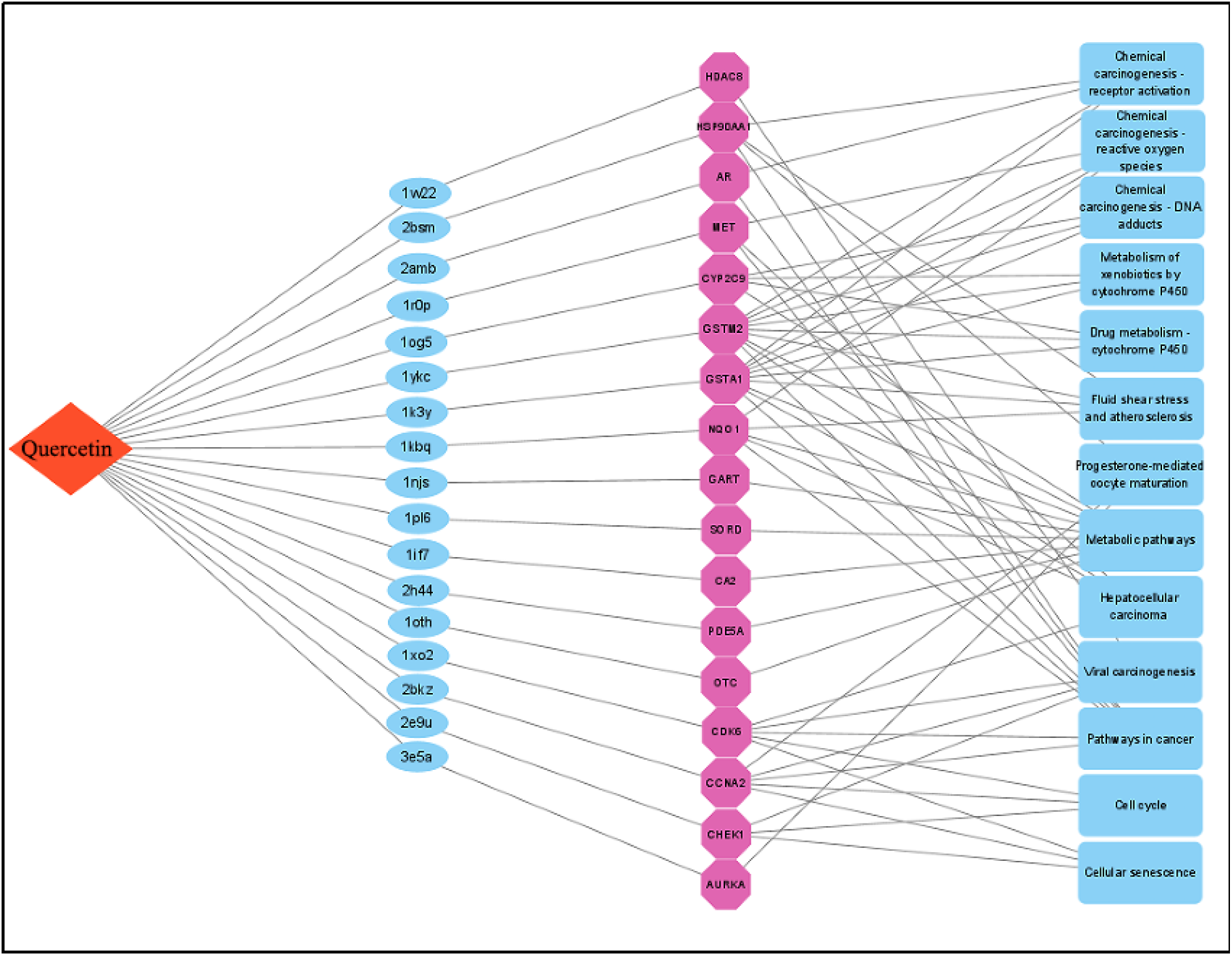
Quercetin-protein-gene-pathway network diagram.

**Figure 8.**
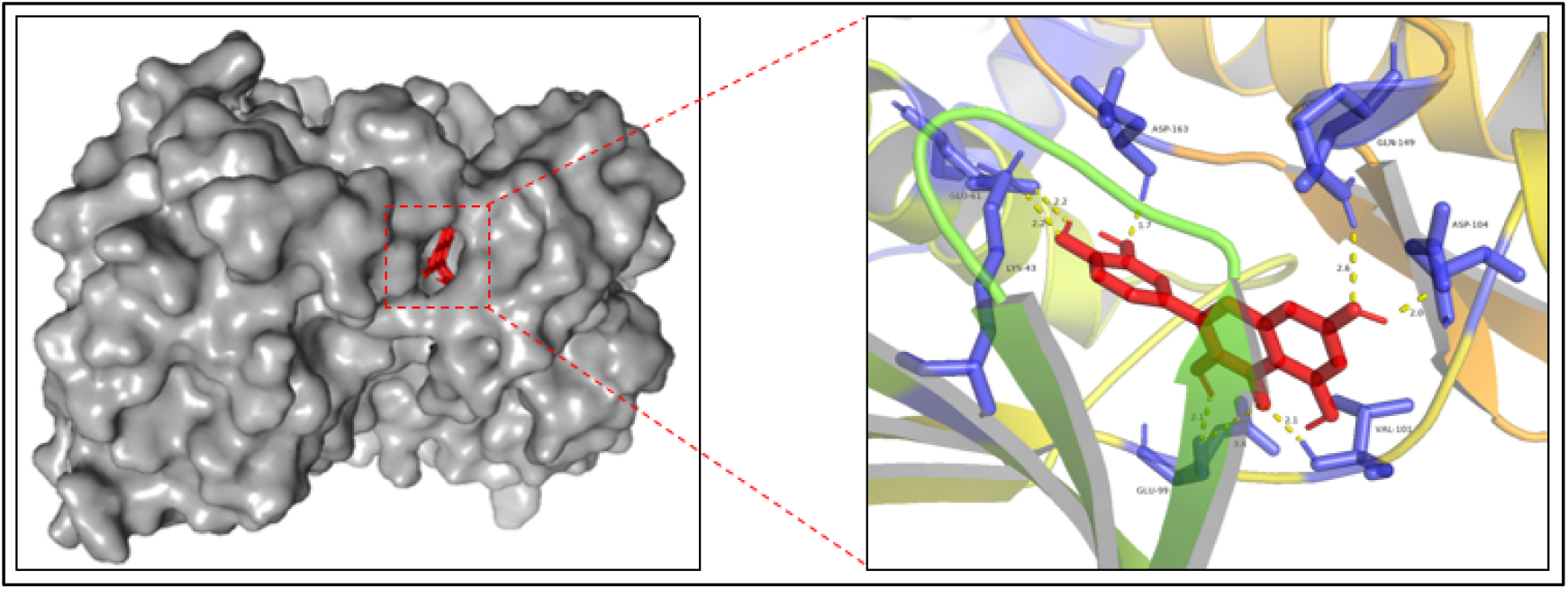
Docking picture of quercetin and 1xo2 protein.

**Figure 9.**
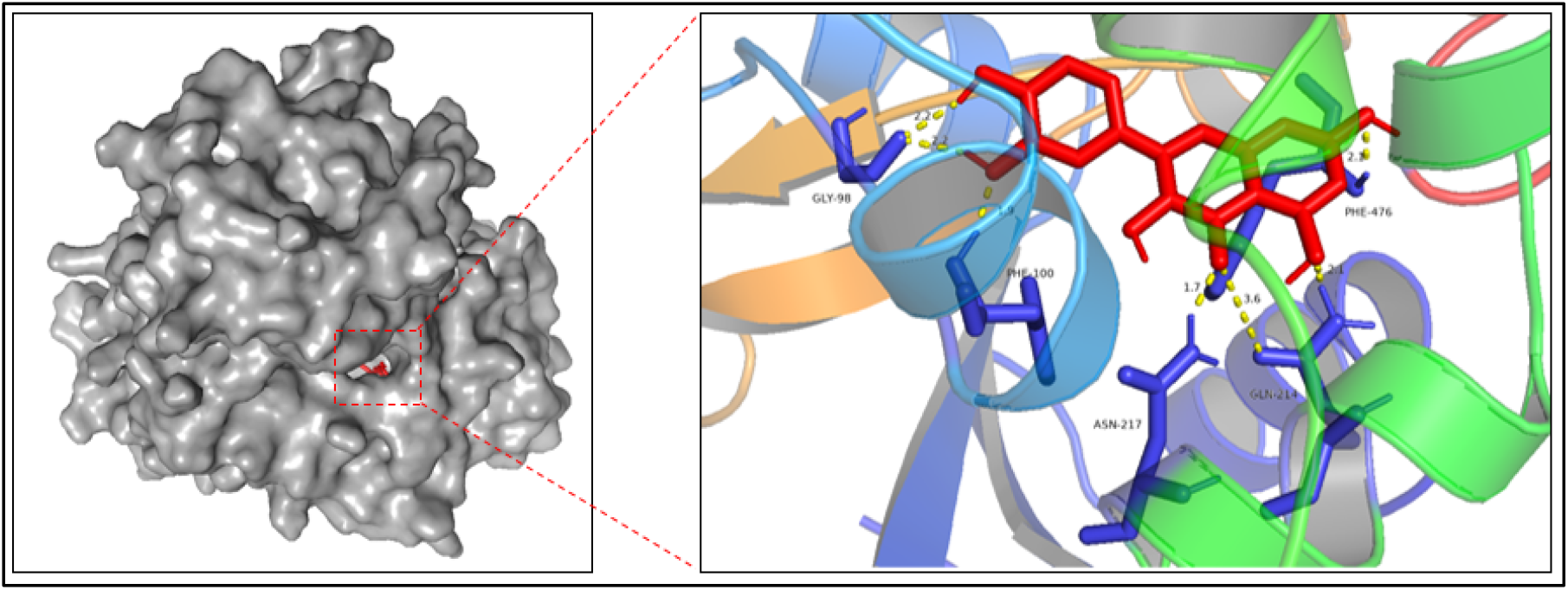
Picture of docking of quercetin with 1og5 protein.

**Figure 10.**
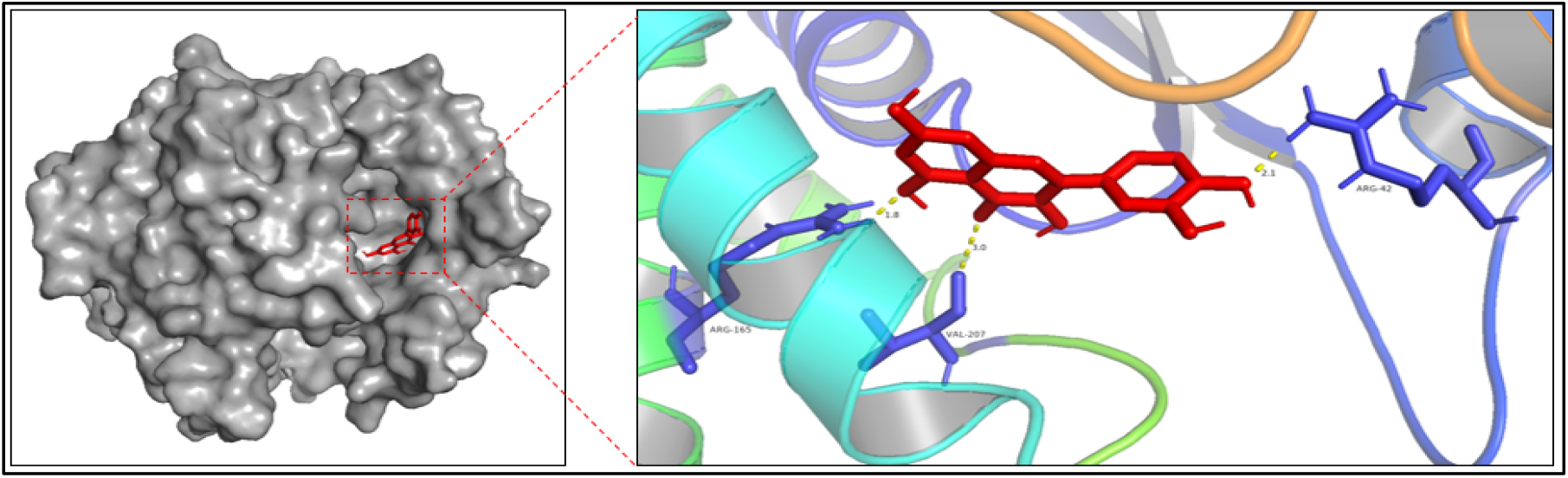
Picture of docking of quercetin with 1ykc protein.

**Figure 11.**
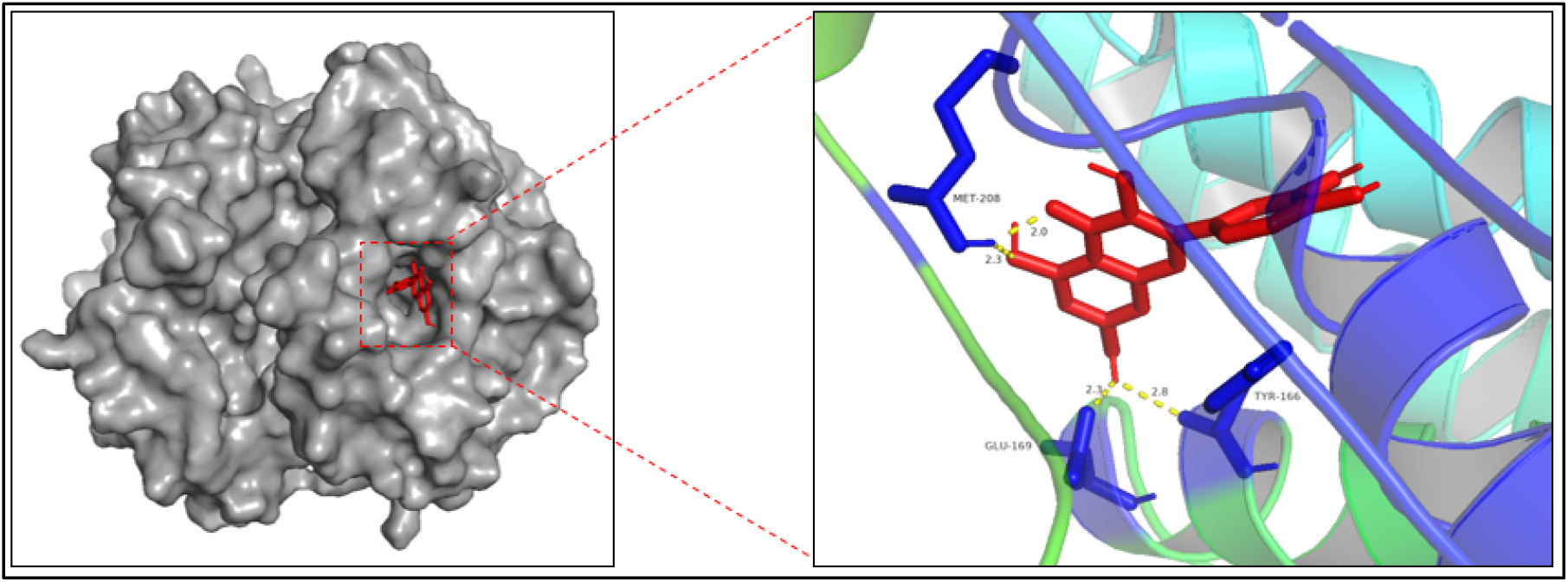
Docking picture of quercetin and 1k3y protein.

**Figure 12.**
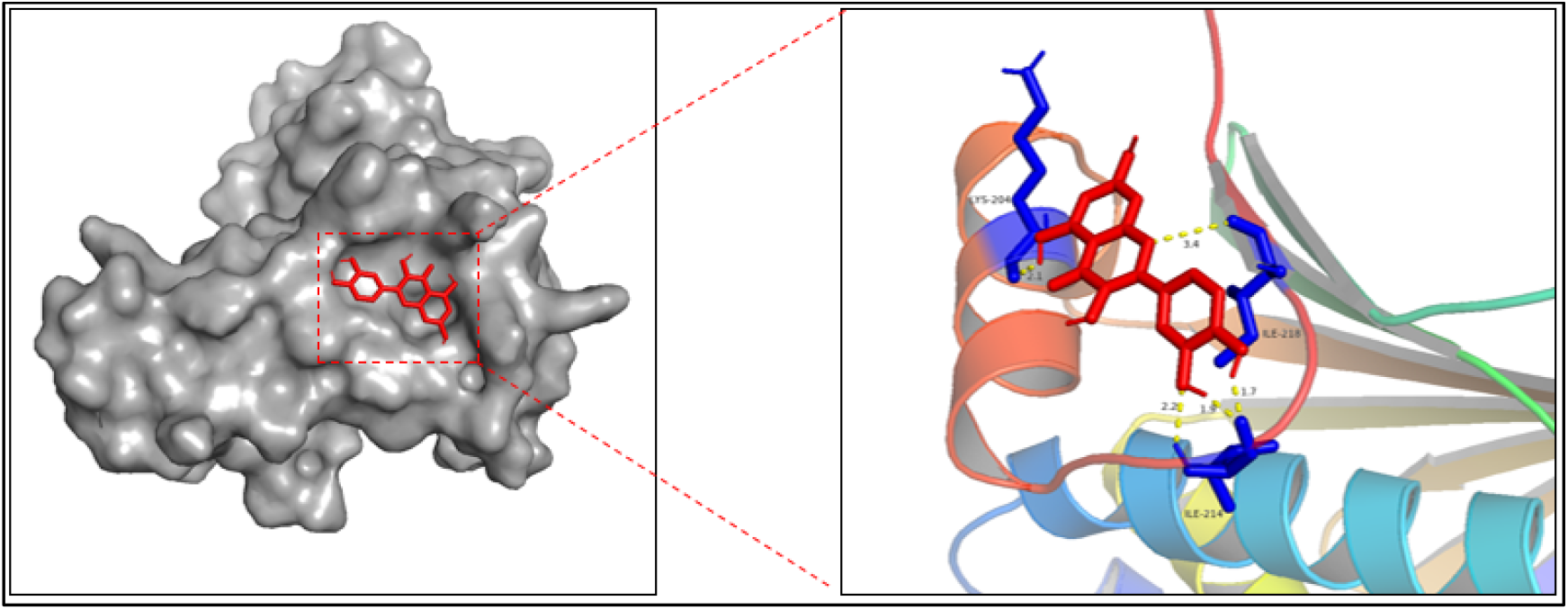
Docking picture of quercetin and 2bsm proteins.

**Figure 13.**
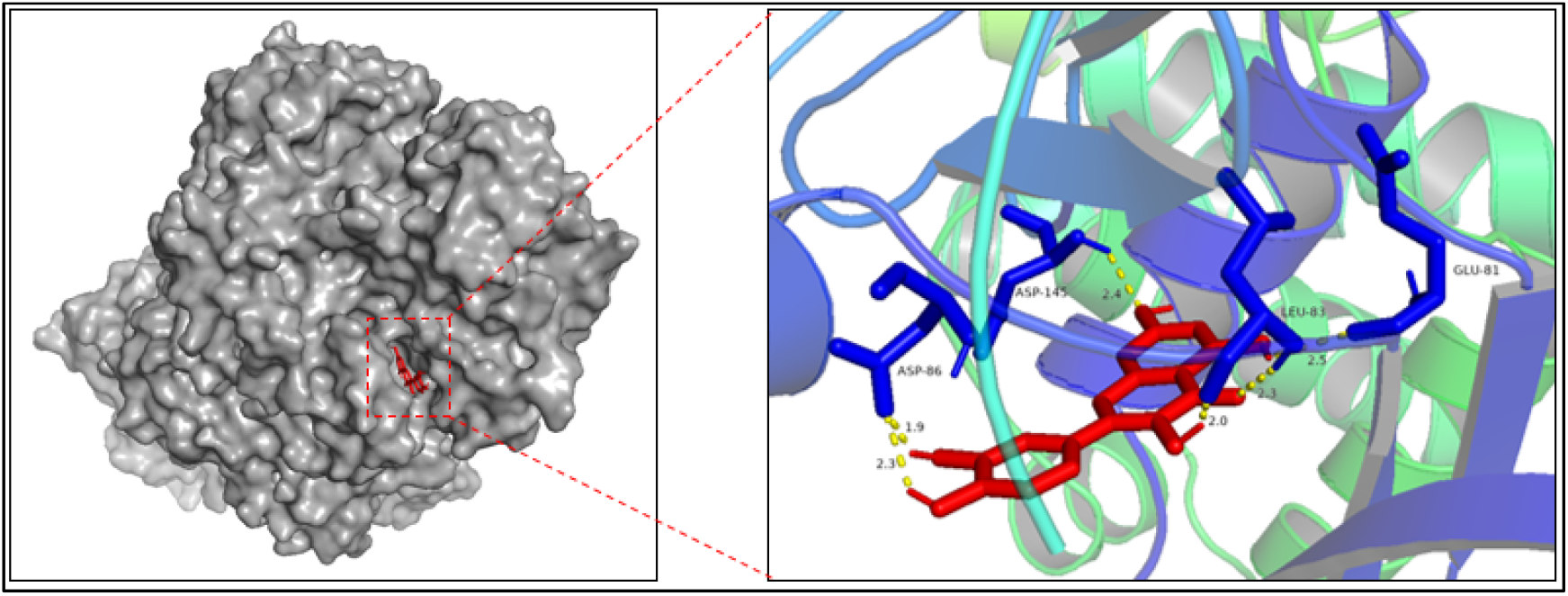
Quercetin and 2 BKZ protein docking images.

**Figure 14.**
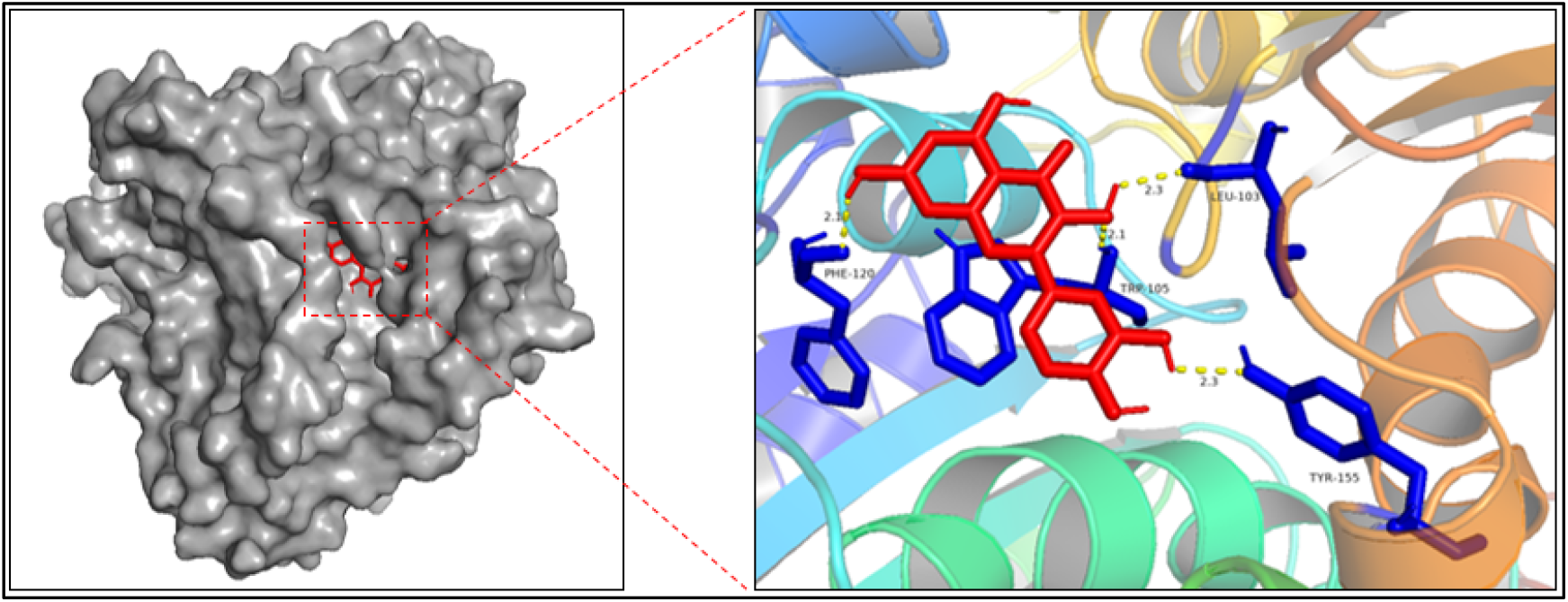
Quercetin and 1 KBQ protein docking images.

### 3.5 Molecular Docking

Selected according to the results of the KEGG GSTM2, GSTA1, NQO1, CCNA2, CDK6, CYP2C9, HSP90AA1 altogether seven target genes, and selected according to the results of the (3.1) seven target genes corresponding protein 1 KBQ k3yykc, 1, 1, 2, BKZ BSM og5 xo2, 1, 1, 2. Using ChemDraw 20.0 and pymol software to pretreatment of protein and quercetin, use AutoDock software for molecular docking, with pymol software for visual processing.

Quercetin and 1 ykc, LYS, GLU, ASP - 163-61-43, ASP, GLU - 99, VAL - 101-104, GLN - 149 form hydrogen bond; With 1 og5 GLY - 98, PHE - 100, the ASN - 217, GLN - 214, PHE - 476 form hydrogen bond; With 1 ykc ARG - 165, VAL - 207, ARG - 42 form hydrogen bond; It formed hydrogen bonds with MET-208, GLU-169, and TYR-166 in 1k3y. LYS-204, ILE-218, and ILE-214 in 2bsm formed hydrogen bonds. It also formed hydrogen bonds with ASP-86, ASP-145, LEU-83 and GLU-81 in 2bkz. And PHE-120, TRP-105, TYR-155 and LEU-103 in 1kbq. The binding energies of quercetin with 1ykc, 1og5, 1ykc, 1k3y, 2bsm, 2bkz, and 1kbq are all less than 5 kJ/mol, demonstrating excellent stability(Table 1).

**Table 1.**
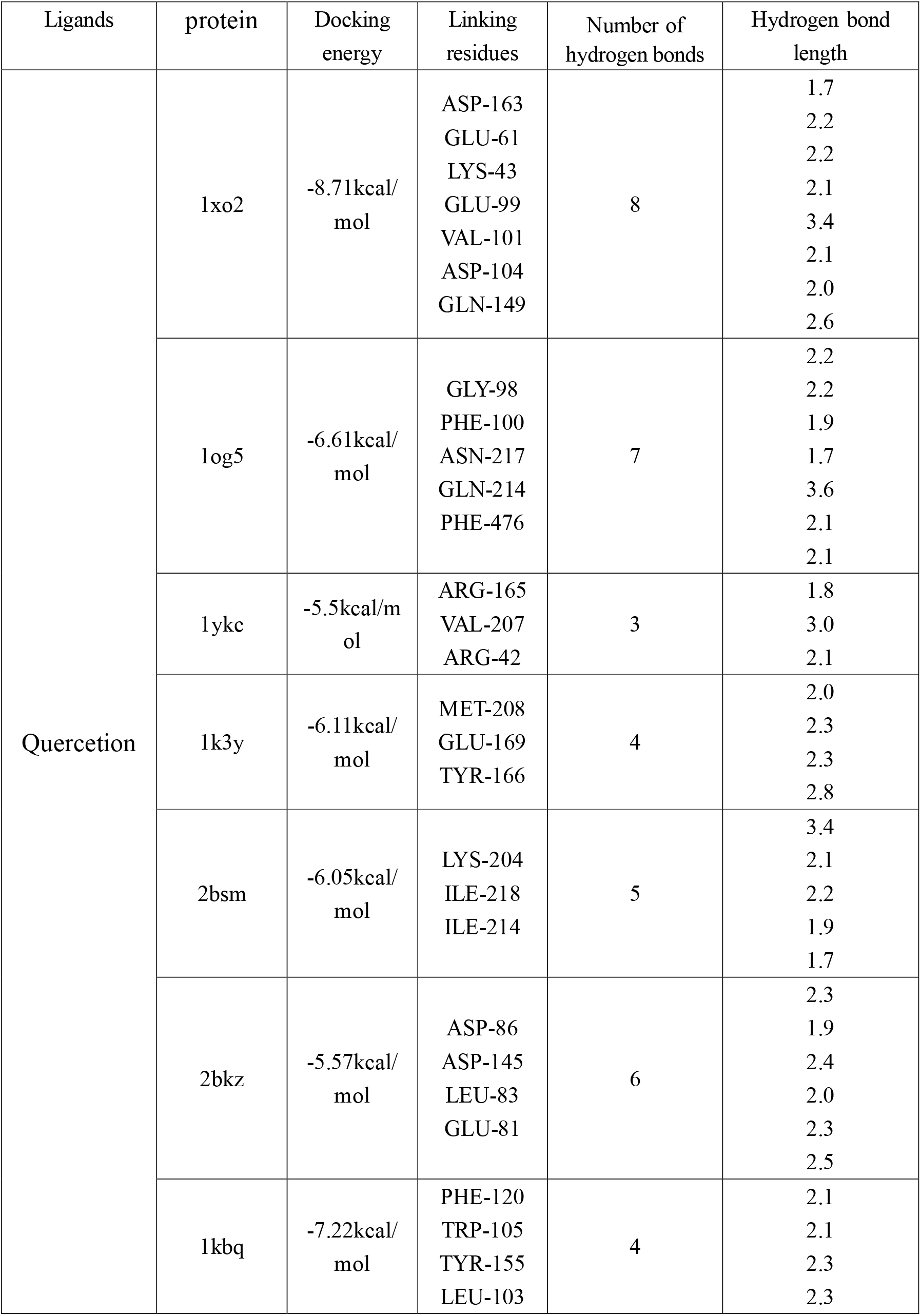
Quercetin and docking protein data.

## 4. Conclusion

In this study, we explored the potential therapeutic targets of quercetin in the treatment of colorectal cancer by network pharmacology, and screened 24 potential target genes. According to the target genes, intracellular steroid hormone receptor signaling pathway, cell division, G2/M transition of mitotic cell were screened Cycle, the apoptotic process, xenobiotic metabolic process, glutathione derivative biosynthetic process, regulation of G2/M transition of mitotic cell cycle, signal transduction, Pathways in cancer, Hepatocellular carcinoma and other signaling pathways, Reflects the role of quercetin multiple targets and pathways characteristics, preliminary explored the quercetin treatment of colorectal cancer biology way; Furthermore, molecular docking technology was used to verify the possibility of predicting the target in the treatment of colorectal cancer to some extent.

## 5. Discussion

According to the results of the seventh National census released by the National Bureau of Statistics in May 2021, the proportion of the population aged 60 and above was 18.70 percent, an increase of 5.44 percentage points compared with the sixth census in 2010. With the trend of aging population becoming more severe, the proportion of elderly colorectal cancer patients will also become more obvious, which will have certain impacts on the medical system of society[18,19].

The study for subsequent quercetin treatment of colorectal cancer provides a certain reference direction. However, this study is only a prediction analysis, and the reliability of the results needs to be further verified by many follow-up experiments.

## Authors’ contributions

Yuanyuan Qian participated in the design of the study and review. Zhaojunli Wang and Jiancheng Ji drafted the manuscript and participated in the data collection and analysis. Wenlong Fu participated in the data collection All authors contributed to the article and approved the submitted version.

## Data availability statement

The datasets used and analyzed during the current study are available from the corresponding author upon reasonable request.

## Acknowledgements

Not applicable.

